# Identification and characterisation of the cryptic Golgi Apparatus in *Naegleria gruberi*

**DOI:** 10.1101/221721

**Authors:** Emily K. Herman, Lyto Yiangou, Diego M. Cantoni, Christopher N. Miller, Francine Marciano-Cabral, Erin Anthonyrajah, Joel B. Dacks, Anastasios D. Tsaousis

**Affiliations:** Department of Cell Biology, University of Alberta, Edmonton, Alberta, Canada.; Laboratory of Molecular and Evolutionary Parasitology, RAPID group, School of Biosciences, University of Kent, Canterbury, Kent, UK; Department of Microbiology and Immunology, Virginia Commonwealth University, School of Medicine, 1101 E. Marshall St, Richmond, VA 23298-0678, USA.

**Keywords:** COPI, protist, evolutionary cell biology, membrane-trafficking, Brefeldin A, dictyosome

## Abstract

Although the Golgi apparatus has a conserved morphology of flattened stacked cisternae in the vast majority of eukaryotes, the organelle has lost the stacked organization in several eukaryotic lineages raising the question of what range of morphologies is possible for the Golgi. In order to understand this range of organellar diversity, it is necessary to characterise the Golgi in many different lineages. Here we identify the Golgi apparatus in *Naegleria*, the first description of an unstacked Golgi organelle in a non-parasitic eukaryote, other than fungi. We provide a comprehensive list of Golgi-associated membrane trafficking genes encoded in two separate species of *Naegleria* and transcriptomic support to show that nearly all are expressed in mouse-passaged *N. fowleri* cells. We then study distribution of the Golgi marker *Ng*COPB by fluorescence, identifying membranous structures that can be disrupted by Brefeldin A treatment consistent with Golgi localisation. Confocal and immuno-electron microscopy revealed that *Ng*COPB is localized to membranous structures consistent with tubules. Our data not only identify the Golgi organelle for the first time in this major eukaryotic lineage, but also provide the rare example of a tubular form of the organelle representing an important sampling point for the comparative understanding of Golgi organellar diversity.

## Introduction

In mammalian cells and many diverse eukaryotes, the Golgi apparatus appears as a stack of flattened membranes, or cisternae, inside which proteins are modified via glycosylation and transported to the plasma membrane or to endolysosomal organelles. In mammalian cells, disrupting the stacked structure of the Golgi leads to numerous defects in these processes, and Golgi fragmentation is observed in autoimmune diseases, cancer, Huntington’s, Parkinson’s, and Alzheimer’s diseases (Zhang and Wang, 2016 *inter alia*). However, a few examples of eukaryotes with unstacked Golgi are found across the evolutionary tree, and include the budding yeast *Saccharomyces cerevisiae* (Preuss et al. 1992), as well as parasitic taxa such as *Plasmodium falciparum, Entamoeba histolytica*, and *Giardia intestinalis* (Mowbrey and Dacks, 2009 *inter alia*).

Many of these organisms were initially thought not to possess Golgi and even to have speciated away from the rest of eukaryotes prior to the origin of the organelle. We now know that this is not the case, based on various lines of evidence. The question therefore becomes, given the conservation of stacked Golgi morphology in the vast majority of eukaryotes, what other structural diversity exists in Golgi organelles? In *S. cerevisiae*, the Golgi compartments are dispersed in the cytoplasm, appearing as punctae using immunofluorescent staining of Golgi markers (Wooding & Pelham 1998). *E. histolytica* was originally thought to not have a Golgi, but an ultrastructural study showed that this was an artefact of the fixation process for transmission electron microscopy, and presented micrographic evidence of dispersed cisternae in the cell (Chávez-Munguia et al. 2000). Finally, Golgi functions in *G. intestinalis* are stage specific, and are carried out in encystation-specific vesicles that form dispersed compartments (Stefanic et al. 2009). Unlike typical Golgi, they are not steady state organelles, but arise in response to cyst wall material formation in the endoplasmic reticulum (Marti, Regös, et al. 2003; Marti, Li, et al. 2003).

Regardless of morphology, organisms with unstacked Golgi encode the membrane trafficking factors necessary for Golgi function (Mowbrey & Dacks 2009). An extensive set of membrane-trafficking machinery has been characterized as involved in vesicle trafficking events to, from and within the Golgi, with distinct paralogues or complexes acting at these steps. These have been shown to be conserved across eukaryotes (Klute et al. 2011) and have been used as genomic signatures for the presence of Golgi organelles in many of the lineages thought once to lack the organelle. However, further characterization of cryptic Golgi in evolutionarily dispersed linages is necessary to have a better understanding of Golgi organellar evolution and the diversity of form that is possible for this organelle.

*Naegleria gruberi* is a free-living microbial eukaryote that is evolutionarily distant from animals, yeast, and plants. It is commonly found in both aerobic and microaerobic soil and freshwater environments worldwide (De Jonckheere 2002; FULTON 1993; Fulton 1974). The closely related *Naegleria fowleri* is an opportunistic neuropathogen of humans and animals, killing ~95% of those it infects within two weeks (Carter 1970). In 2010, the *N. gruberi* genome was published, revealing a remarkably complex repertoire of cytoskeletal, sexual, signalling, and metabolic components (L. K. Fritz-Laylin et al. 2010), as well as a highly complete membrane trafficking system (MTS). Naegleria is a member of the supergroup Excavata, which also contains the trypanosomatids, *Trichomonas vaginalis*, and *Giardia intestinalis*, of parasitological importance, and the rodent gut commensal *Monocercomonoides* sp., which is both anaerobic and amitochondriate. Thus, *N. gruberi* remains one of the few free-living excavates with a complete and publicly available genome, making it a key sampling point for studying eukaryotic evolution, and potentially a useful model system for studying eukaryotic cell biology outside of the animals, yeast, and plants. One distinctive cellular feature of *Naegleria* is that it lacks a visibly identifiable Golgi organelle. This is in fact a diagnostic feature of the larger taxonomic group to which *Naegleria* belongs, the heteroloboseans (Adl et al. 2012) and has been since its inception (Page & Blanton 1985).

Despite some proposals for membranous structures as putative homologues of the Golgi (Stevens et al. 1978), the only evidence supporting the presence of the organelle in *Naegleria* has been the bioinformatically predicted Golgi-associated proteins, identified in the genome project (L. K. Fritz-Laylin et al. 2010). We here address the identification and visualisation of the Golgi structure in *Naegleria* using a multidisciplinary approach and present the first molecular and cellular evidence for the presence of a punctate Golgi in *N. gruberi*, distinct from other endomembrane organelles.

## Methods

### Comparative Genomics

For all Golgi-associated MTS genes that were not identified by Fritz-Laylin et al. 2010, the functionally characterized human orthologue was used as a BLASTP or TBLASTN query (Altschul et al. 1997) to search the *N. gruberi* NEG-M (Joint Genome Institute, http://genome.jgi.doe.gov/Naegr1/Naegr1.home.html) predicted proteome, EST cluster consensi, and scaffolds. *N. gruberi* sequences retrieved with an E-value of 0.05 or less were used to reciprocally BLAST the *H. sapiens* and non-redundant protein databases (NCBI, https://www.ncbi.nlm.nih.gov/). To be considered true orthologues, they must retrieve the initial query or a clear orthologue with an E-value less than the 0.05 cut-off. Comparative genomics searches of the *N. fowleri* transcriptome were performed using the *N. gruberi* Golgi-associated MTS sequences as queries, or the human orthologue in cases where the *N. gruberi* sequence could not be identified, with the same criteria as above.

### Phylogenetics

Bayesian and Maximum-Likelihood phylogenetics analyses were performed to assign orthology to sequences in the Qa- and Qb-SNARE families, and the Qc-SNARE subfamily including Syntaxin 6, Syntaxin 8, and Syntaxin 10. Phylogenetically characterized sequences from *H. sapiens*, *Arabidopsis thaliana*, and *Trypanosoma brucei* were aligned with *N. gruberi* and *N. fowleri* sequences identified in this study using MUSCLE v.3.8.31 (Edgar 2004). Alignments were visualized in Mesquite v.3.2 (Maddison and Maddison 2017), and manually masked and trimmed to remove positions of uncertain homology. ProtTest v3.4 (Darriba et al. 2011) was used to determine the best-fit model of sequence evolution, which was LG+G+F for the Qa- and Qb-SNARE alignments, and LG+G for the Syntaxin 6/8/10 alignment. Phylobayes v4.1 (Lartillot et al. 2009) and MrBAYES v3.2.2 (Ronquist et al. 2012) programs were run for Bayesian analysis and RAxML v8.1.3 (Stamatakis 2014) was run for Maximum-Likelihood analysis. Phylobayes was run until the largest discrepancy observed across all bipartitions was less than 0.1 and at least 100 sampling points were achieved, MrBAYES was used to search treespace for a minimum of one million MCMC generations, sampling every 1000 generations, until the average standard deviation of the split frequencies of two independent runs (with two chains each) was less than 0.01. Consensus trees were generated using a burn-in value of 25%, well above the likelihood plateau in each case. RAxML was run with 100 pseudoreplicates.

### Transcriptomics

*N. fowleri* (Ax) V212 were grown axenically at 37°C in Oxoid medium in T75 culture flasks (CLINE et al. 1983). A mouse-passaged strain of V212 was obtained by intranasal inoculation of the amoebae in B6C3F1 mice. Mice were sacrificed when symptoms of infection were evident. The amoebae were harvested from brain tissue and then continuously passaged through mice two times (MP2), four times (MP4) and six times (MP6). Amoebae were subsequently cultured as above for 7-10 days to remove residual brain tissue, and then RNA was extracted and converted to cDNA with the Affymetrix/USB M-MLV (cat 78306) kit using standard protocols. Illumina libraries were constructed using the Nextera Workflow and sequenced in an Illumina MiSeq at the TAGC facility (UAlberta).

Between 2.8-4.0 million paired-end 300bp reads remained after pre-processing with Trimmomatic v0.36 (Bolger et al. 2014) using the arguments SLIDINGWINDOW:50:30 TRAILING:20. Reads were aligned to an unpublished genome of *N. fowleri* strain V212 produced as part of an on-going project (unpublished) using the program Tophat v2.0.10 (Trapnell et al. 2009). Transcripts were assembled for each condition using Cufflinks v2.1.1 and then merged with Cuffmerge (Trapnell et al. 2010). Cuffdiff v2.1.1 (Trapnell et al. 2013) was then used to map the reads to the merged transcripts and determine the relative expression. For some Golgi-associated MTS genes, the merged transcript appeared to be partial or incorrectly fused with another gene product, based on comparison with the *N. gruberi* homologues. To more accurately assess the expression of these transcripts, the transcript boundaries were manually modified to correct for this before mapping the reads. All newly generated *N. fowleri* gene sequences have been deposited in Genbank as Accession (XXXXX-YYYYY).

### *Naegleria* cell culturing

*Naegleria gruberi* strain NEG-M (kindly provided by Lillian Fritz-Laylin) was grown axenically at 28 °C in M7 medium (Fulton 1974). Cells were passaged every 3 – 5 days depending on their density.

### RNA extraction

Total RNA extraction of *Naegleria gruberi*’s trophozoites was performed using RNAeasy Midi Kit (Qiagen) according to the manufacturer’s protocol. cDNA was amplified according to the manufacturer’s guidelines using the SuperScript III RT Reaction (Invitrogen).

### Generation of antisera to N. gruberi proteins

The entire *N. gruberi* NEG-M strain ORFs of COPB (XP_002673194) and Sec31 (XM_002669379) were PCR amplified from the cDNA using the primers shown in Table S1 and subsequently cloned into pET16 or pET30b (Novagen), sequenced and transformed into *Escherichia coli* BL21(DE3) PLyS cells. The expressed proteins were purified using a Ni-NTA column under native conditions. The proteins were further purified by gel electrophoresis and 4 mg of each purified proteins were used to make chicken and rat polyclonal antisera, respectively (two animals per protein; Davids Biotechnologie GmbH; Germany).

### Cell Fractionation of Naegleria

*Naegleria gruberi* cellular fractions were achieved using differential centrifugation of the cell homogenate (passed five times through a gauge 33 hypodermic needle). All steps were carried out at 4 °C and in the presence of the protease inhibitors (Complete Mini EDTA-free cocktail tablets, Roche). To separate cellular fractions, the cells were centrifuged at 1,000 × g for 10 min, and washed and resuspended in the buffer (250 mM sucrose and 20 mM MOPS, pH 7.4). The homogenate was centrifuged at 1,000 × g for 10 min to remove unbroken cells. The supernatant was then centrifuged at 3,000 × g for 15 mins to collect the pellet that contained the nuclei. The supernatant was carefully collected and centrifuged at 7,000 × g for 30 minutes to discard the mitochondrial fraction (Tsaousis et al. 2014). The membrane fraction was centrifuged at 20,000 × g for 3 hours and the supernatant was used as the cytosolic fraction. The separated fractions were quantified using a Bradford assay (Bio-Rad) and then were analyzed via western blot analysis.

### Western blotting

Western blots of total protein extracts from *N. gruberi* trophozoite cells were incubated with the chicken anti-Cop1 (1:200) and rat anti- Sec-31 (1:200) antisera, followed by secondary anti-chicken and anti-rat antibodies respectively conjugated to peroxidase (Sigma). The blots were developed using the ECL protocol (Amersham) and visualized using the Syngene G:BOX XT4 machine on the GeneSys software.

### Immunofluorescence microscopy

*Naegleria gruberi* cells were seeded at 30,000 cells/ml on NUNC LabTek Chamber slides prior to the experiment and grown for 24 hours. Cells were incubated for 20 min with the ER-tracker blue-white DPX marker (Molecular Probes) and then fixed using 2% formaldehyde followed by permeabilization with 0.1% Triton-X in 1X PBS. After blocking for 1 hour in 3% BSA – 1X PBS, the cells were probed with the chicken anti-COPI (1:250) and rat anti- Sec (1:250) antisera. Secondary Alexa Fluor 488 goat anti-chicken IgG (H-L), Alexa Fluor 488 chicken anti-rat IgG and Alexa Fluor 594 donkey anti-rat IgG (Molecular Probes) were used in dilution of 1:1000 to the corresponding primary antibodies. Cells were mounted with DAPI-containing anti-fade mounting reagent (Vectashield), and observed under an Olympus IX81 fluorescence microscope and a laser scanning Zeiss LSM 880 confocal microscope. Images were collected using Micromanager 1.4 software for fluorescence and Zeiss Zen software for confocal microscope and processed with ImageJ.

### Fixation of cells for electron microscopy and immuno-gold labelling

Aspirated cultures of *N. gruberi* were fixed for 1 hour in freshly prepared PBS solution containing either 2.5% Glutaraldehyde or 4% Formaldehyde, for Contrast Transmission Electron Microscopy (CTEM) or Immuno-Electron Microscopy (IEM) respectively. Both sample preparations were then washed several times with PBS. CTEM samples were stained and postfixed with 1% Osmium. Both sample preparations were then dehydrated through an ethanol series (30%, 50%, 70%, 90% and three times in 100%). The ethanol was then aspirated and replaced with the appropriate resin mixture. CTEM samples were suspended in Propylene Oxide (PO, Agar Scientific) and mixed thoroughly by rotor, the PO was then replaced by a 50:50 mix of PO and Agar Scientific Low Viscosity (LV) Resin and mixed again. The PO:LV resin mix was then replaced by 100% LV resin. IEM samples were similarly suspended in LR white Resin (Agar Scientific). Resin permeation was aided by placing the samples in a vacuum for 2 minutes. The resin was then aspirated and replaced with fresh resin and the samples transferred into Beem (for CTEM, Agar Scientific) or Gelatin (for IEM, Agar Scientific) capsules and hardened for 15 hours in a pre-warmed 60°C Oven. The hardened blocks were then polished and subsequently sectioned by ultra-microtome at a thickness of 70 μm, then placed on 50 mesh, 3 mm copper or gold EM grids for CTEM or IEM respectively at approximately 5 sections per grid. Immuno-staining of the IEM grids was performed in humidifying chambers. Blocking of the samples was achieved via a 1 hour incubation in 2% Bovine Serum Albumin in PBS-0.05%Tween. Primary antibody binding was performed by 15 hour incubations with the primary antibody, at three dilutions (1:10, 1:50 and 1:100) at 8°C. The IEM grids were subsequently incubated for 30 minutes at room temperature, with the corresponding gold-conjugated secondary antibodies. Counter-staining of both CTEM and IEM sample grids was achieved via a 15 minutes’ incubation with 4.5% Uranyl acetate in PBS and a 2-minute incubation in Reynold’s Lead Citrate.

### Brefeldin A experiments

Prior to each experiment (immunofluorescence microscopy or protein extraction/cell fractionation), *Naegleria* cells were incubated at 28 °C in M7 medium for 3 hours with 10 nM, 100 nM and 1 μM of Brefeldin A (S7046-SEL, Stratech Scientific Ltd).

## Results

### *Naegleria* encode and express Golgi-associated membrane trafficking machinery

*Naegleria*, along with the rest of the heteroloboseans, have been diagnostically described as lacking a visible stacked Golgi (Page & Blanton 1985; Adl et al. 2012). However, sequencing and annotation of the *N. gruberi* genome suggested that it encodes many of the necessary components for Golgi function (L. K. L. K. Fritz-Laylin et al. 2010).

We first sought to expand this list in *N. gruberi*. In cases where homologous sequences could not be identified in Fritz-Laylin *et al*. (L. K. L. K. Fritz-Laylin et al. 2010), we performed additional BLAST searches using functionally characterized human sequences as queries to search the genome and predicted proteome of *N. gruberi* (Table S1). We identified several additional homologues not originally reported in the genome paper, including three members of the COG tethering complex, a nearly complete EARP/GARP complex, and a single Syntaxin 6 orthologue. As the Qa-, Qb-, and Qc- SNAREs are highly paralogous gene families, phylogenetic analyses were performed in order to classify orthologues (Supplementary Figure 1a-c). Therefore, the set of Golgi-associated MTS machinery in *N. gruberi* is even more complete than previously thought (Figure 1, Table S1).

**Figure 1:**
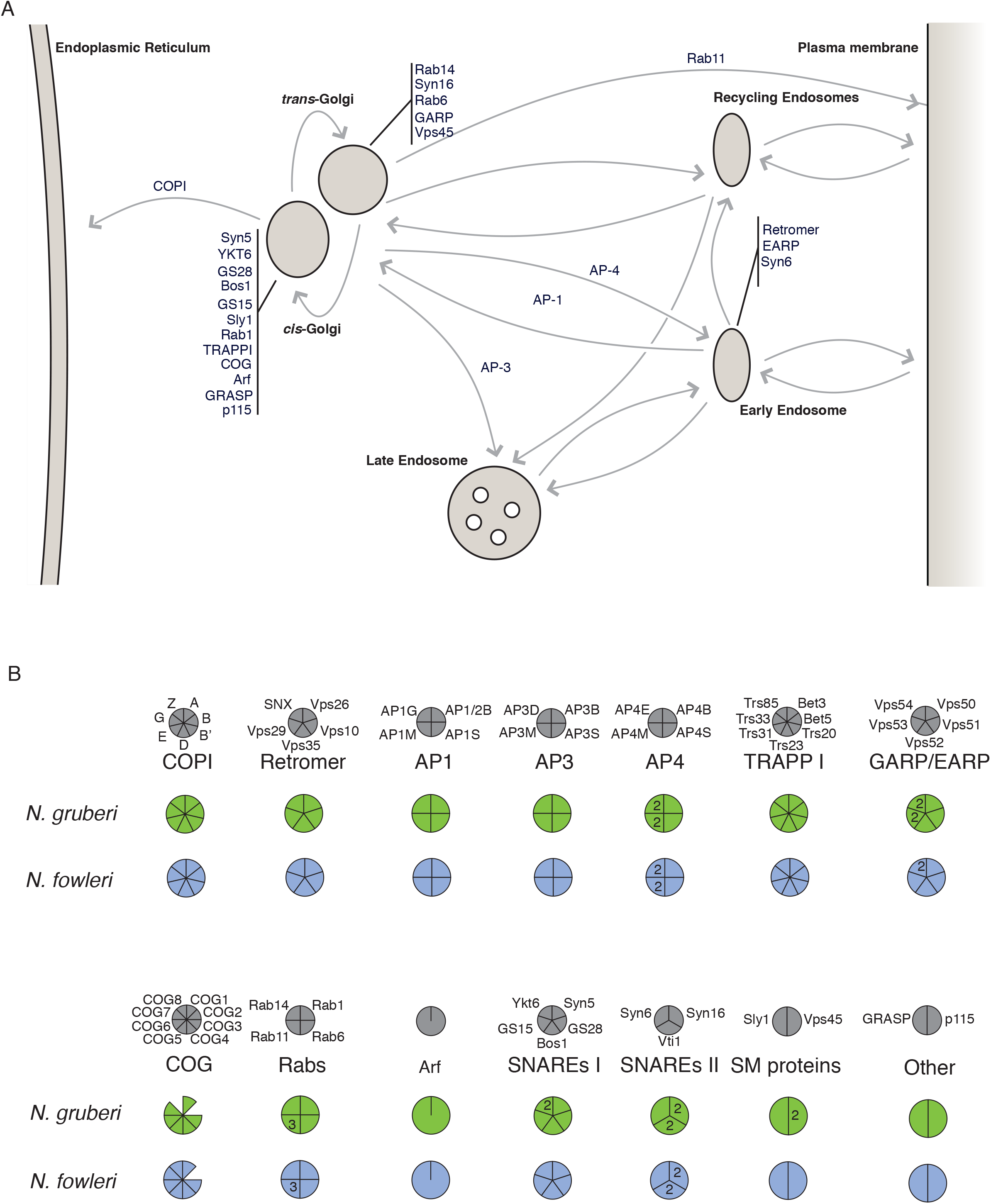
*In silico* prediction of Golgi-associated proteins in *Naegleria*. a. Cartoon illustrating the membrane trafficking pathways in which Golgi-associated proteins function. b. Coulson plot showing the presence of Golgi-associated proteins shown in (a.) in *N. gruberi* and *N. fowleri*. Grey circles represent the complement of Golgi-associated proteins that are generally conserved in eukaryotes. Below, filled segments represent identified homologues, while missing segments indicate that no homologue could be found. Paralogue numbers are shown within the segments.

The presence of these genes in one *Naegleria* species suggests functional relevance. Nonetheless, the possibility remains that the genes may not be translated. In order to determine whether these genes are present in other *Naegleria* species, and more importantly expressed, we generated transcriptomic data from *N. fowleri*. mRNA was extracted from *N. fowleri* grown axenically and mouse-passaged. Homology searching was then performed to identify Golgi-related MTS genes in the resulting transcripts, and the relative expression of these genes under each condition was calculated. We identified expressed transcripts for all 66 Golgi-associated MTS genes in *N. fowleri*, (Table S1). All *N. gruberi* sequences were shown to have an orthologue in *N. fowleri*, with the exception of three *N. gruberi*-specific paralogues (Vps53A, Ykt6B, and Vps45B). Furthermore, *N. fowleri* encodes and expresses two additional members of the COG complex. These results suggest that *Naegleria* not only encodes, but also expresses Golgi trafficking and structural proteins.

### Visualization of the Golgi in N. gruberi

A commonly used marker for Golgi in eukaryotic cells is the COPI complex, which forms vesicle coats for trafficking from the *cis*-Golgi to the ER (Orci et al. 1986; Szul & Sztul 2011) and acts at cis and intermediate Golgi compartments (Papanikou et al. 2015). In order to visualize the *N. gruberi* Golgi using immunofluorescence microscopy, we generated antibodies specific to the beta subunit of the COPI complex (NgCOPB) and to the Sec31 protein of the COPII complex (NgSec31), which is specific to the ER as a comparison point for an endomembrane organelle of the early secretory system. The generated antisera, from chicken and rat respectively, showed high specificity for NgCOPB and NgSec31 in western blots, as they recognized a protein with expected size of 114.5 KDa and 145.7 kDa in the *N. gruberi’s* extracted cell lysates (**Supplementary Figure 2**). Immunofluorescence microscopy of *N. gruberi* cells showed distinct patterns for the two antibodies (**Figure 2**). Consistent with standard ER morphology, the Sec31 antisera showed a network-like localisation around the nucleus of the organism. By contrast, the COPB antisera showed some cytosolic staining, consistent with COPB being a cytosolic/peripheral membrane protein, but strikingly, we see punctuated and tubular localisation around the cell. Some apparent overlap was observed as expected for organelles that span the breadth of the cells. However, clear areas of non-overlap were seen consistent with these structures being discrete organelles.

**Figure 2:**
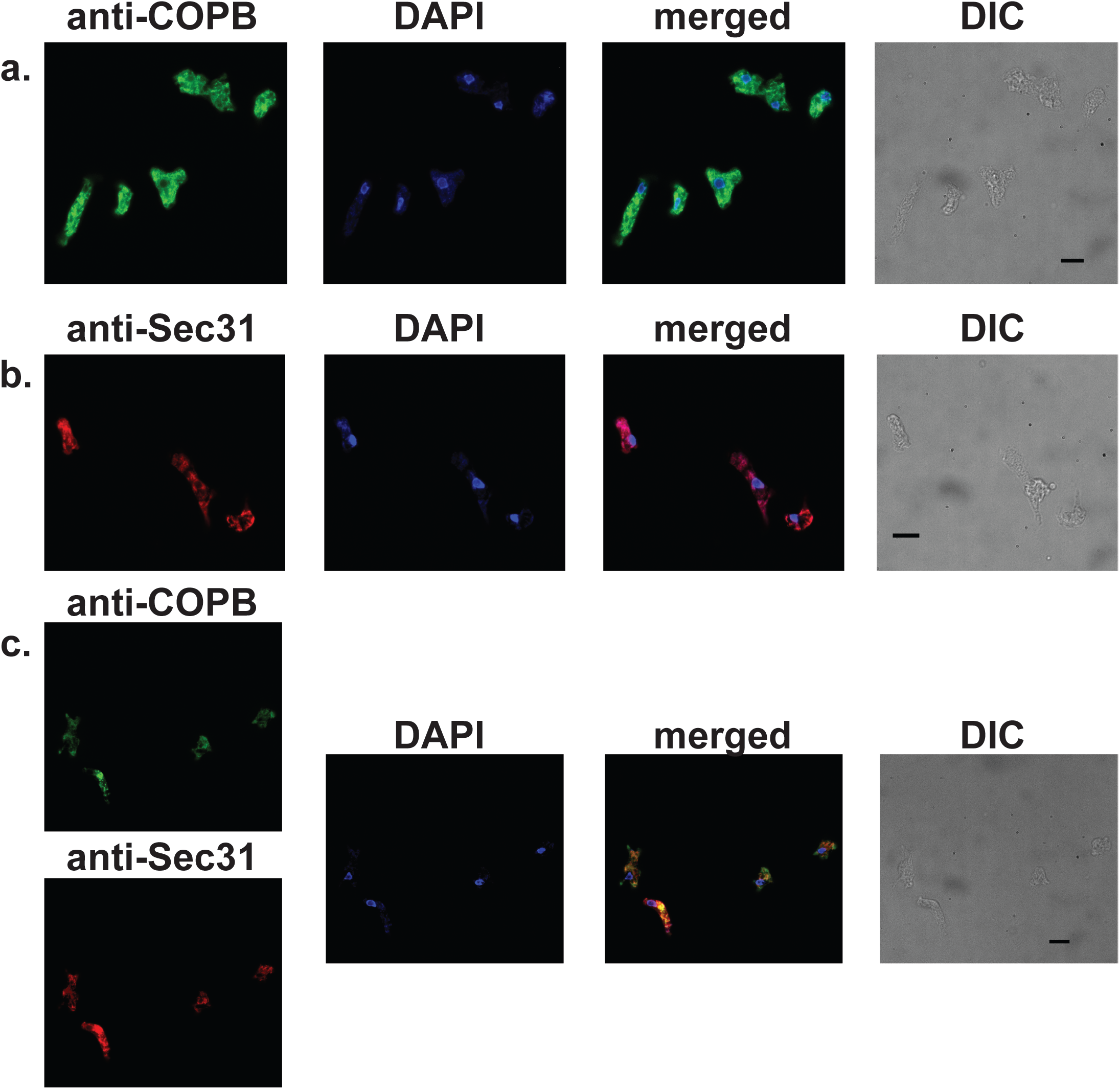
Indirect Fluorescence Assay of *Naegleria* cells. **a.** Cellular localisation of COPI in *N. gruberi* cells. Chicken anti-*N.gruberi* COPB antiserum (1:250; green) shows a discrete localisation in the cells, while DAPI stains the *Naegleria* nucleus and mitochondrial DNA. DIC shows the differential interference contrast image of the cells used for immunofluorescence. Scale bar: 10 μm. **b.** Cellular localisation of Sec31 in *N. gruberi* cells. Rat anti-*N.gruberi* Sec31 antiserum (1:250; red) shows a discrete localisation in the cells, while DAPI stains the *Naegleria* nucleus and mitochondrial DNA. DIC shows the differential interference contrast image of the cells used for immunofluorescence. Scale bar: 10 μm. **c.** Cellular localisation of COPI and Sec31 in *N. gruberi* cells. Chicken anti-*N.gruberi* COPB antiserum (green) shows a discrete localisation in the cells that is not co-localised with the rat anti-*N.gruberi* Sec31 antiserum (red), while DAPI stains the *Naegleria* nucleus and mitochondrial DNA. DIC shows the differential interference contrast image of the cells used for immunofluorescence. Scale bar: 10 μm.

### Brefeldin A treatment disrupts COPB localisation

To further assess whether the COPB antisera was marking Golgi organelles, we tested whether the observed discrete localisation is inhibited by Brefeldin A (BFA), a fungal metabolite that rapidly and reversibly inhibits transport of secretory proteins resulting in relocation of Golgi resident proteins to the ER and COPB to the cytoplasm (Klausner et al. 1992). Sub-cellular fractionation of membrane constituents showed that treatment with BFA at 10 nM, 100 nM and 1 μm for 3 hours shifted the COPB intensity from the internal membrane to the cytosolic fraction (Figure 3a). Consistent with the previous results, immunofluorescence microscopy demonstrated that the punctuated/tubular localisation disappears as concentrations of BFA increase (Figure 3b).

**Figure 3:**
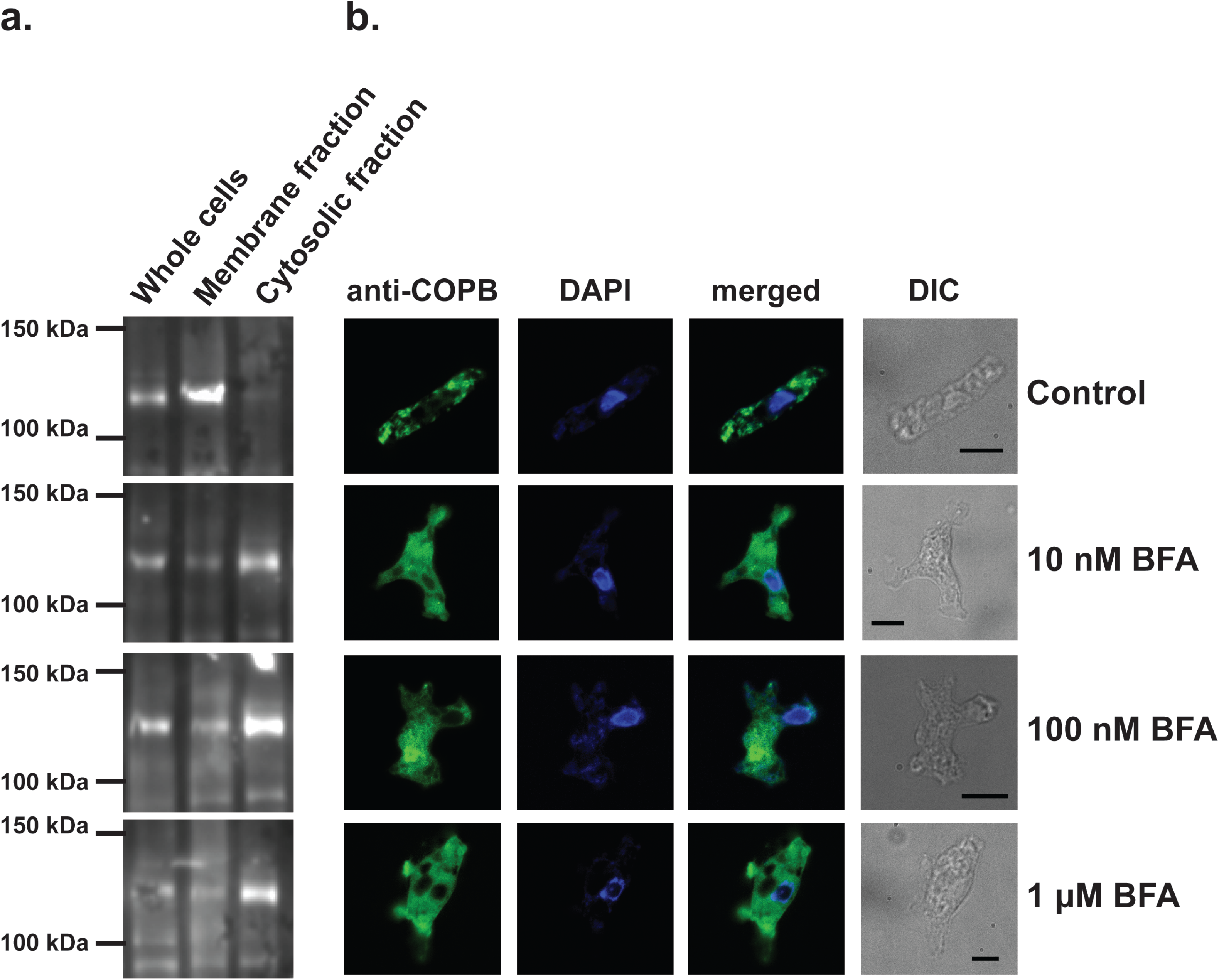
Brefeldin A changes the localisation of COPB in *Naegleria*. **a.** Western blot of whole cells, membrane and cytosolic fraction using the anti-COPB antisera, in the presence of various concentrations or absence of BFA. With the increasing concentrations of BFA from 10 nM to 1 μM after 3 hours of incubation, COPB intensity increases in the cytosolic fraction, while it decreases in the membrane fractions. For each experiment, we have used the same initial number of *Naegleria* cells, and after a Bradford assay, the same amount of protein was loaded in the gel. **b.** Indirect Fluorescence Assay of *Naegleria* cells after BFA treatment. Chicken anti-*N.gruberi* COPB antiserum (1:200; green) shows localisation of COPI in the cells, while DAPI stains the *Naegleria* nuclei and mitochondrial DNA. Same as with the western blot experiments, with the increasing concentrations of BFA from 10 nM to 1 μM after 3 hours of incubation, COPB intensity increases in the cytosol, while the membranous localisation is disappearing. DIC shows the differential interference contrast image of the cells used for immunofluorescence. Scale bar: 10 μm

### The Naegleria Golgi appears as a tubular organelle

We performed localisation experiments captured with confocal microscopy, in order to better assess the ultrastructure of the *Naegleria* Golgi. Using the same antibody concentrations as before, we captured the localisation of COPB in various cells. The 3D rendering of several sections per cell, demonstrate a discrete tubular organisation (**Figure 4a-c, Supplementary Figure 3; Supplementary Movies 1-3**), which was validated in 24 independent experiments using various antibody concentrations. This localisation pattern disappeared, when the cells were incubated with various concentration of BFA for 3 hours: COPB was clearly localised in the cytosol of the *Naegleria* (**Figure 4d-e, Supplementary Figure 4; Supplementary Movies 4-5**). Both localisation patterns were unlike the one seen for Sec31 (**Supplementary Figure 5**).

**Figure 4:**
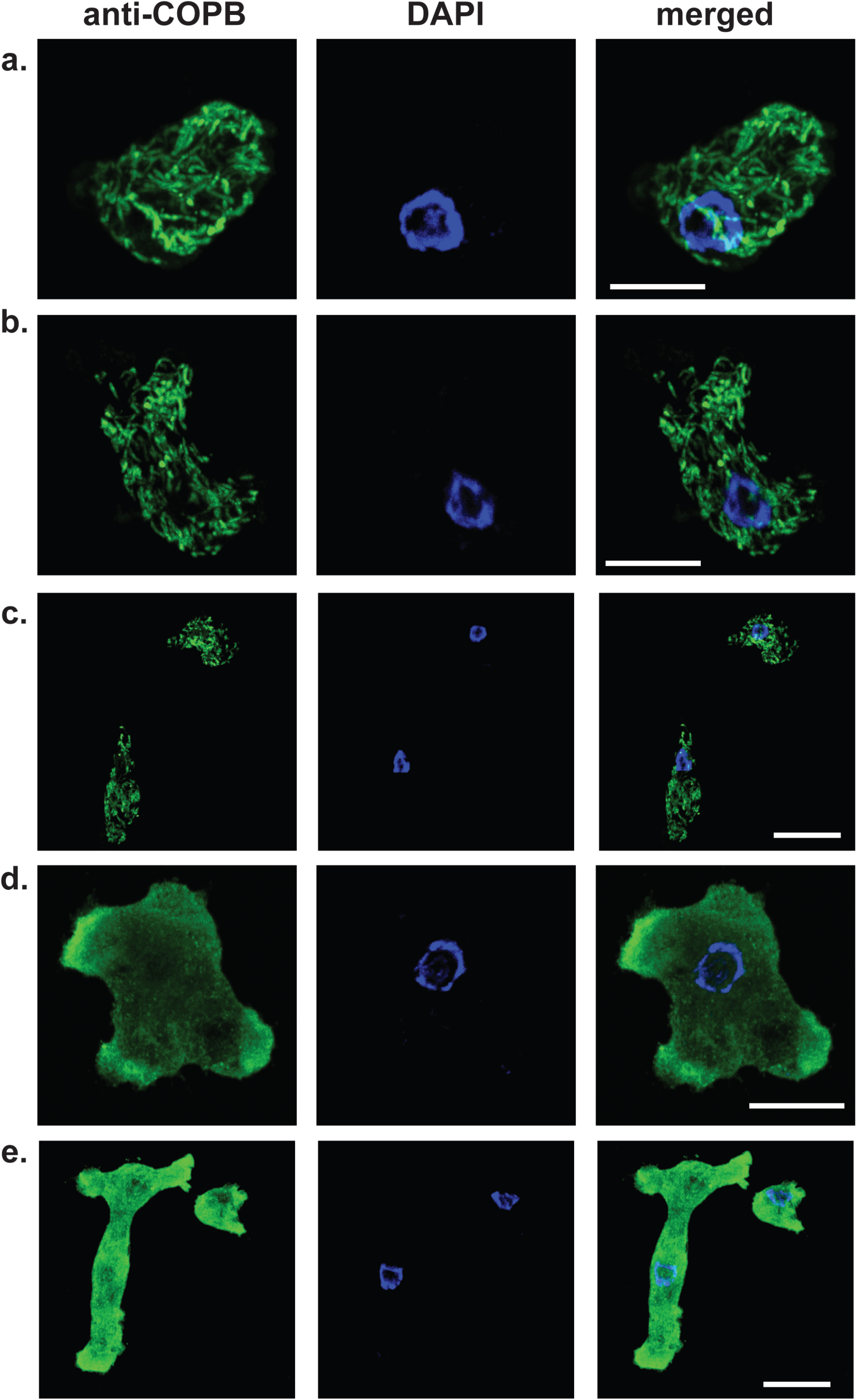
Confocal microscopy of COPI localisation in individual *N. gruberi* cells. **a.** Cellular localisation of COPI in a *N. gruberi* cell. Chicken anti-*N.gruberi* COPB antiserum (1:250; green) shows a tubular localisation in the cell, while DAPI stain the *Naegleria* nucleus and mitochondrial DNA. The image is a result of a 3D rendering of 28 individual, 0.284 μm thick sections, overlapping with a final representative thickness of 7.95 μm. Different angles of the same image can been found in **Supplementary Figure 3a**. **b.** Cellular localisation of COPI in a *N. gruberi* cell. Chicken anti-*N.gruberi* COPB antiserum (1:250; green) shows a tubular localisation in the cell, while DAPI stains the *Naegleria* nucleus and mitochondrial DNA. The image is a result of a 3D rendering of 16 individual, 0.295 μm thick sections, overlapping with a final representative thickness of 4.624 μm. Different angles of the same image can been found in **Supplementary Figure 3b**. **c.** Cellular localisation of COPI in two *N. gruberi* cells. Chicken anti-*N.gruberi* COPB antiserum (1:250; green) shows a tubular localisation in the cell, while DAPI stains the *Naegleria* nuclei and mitochondrial DNA. The image is a result of a 3D rendering of 21 individual, 0.284 μm thick sections, overlapping with a final representative thickness of 5.96 μm. Different angles of the same image can been found in **Supplementary Figure 3c**. **d.** Cellular localisation of COPI in a *N. gruberi* cell. Chicken anti-*N.gruberi* COPB antiserum (1:250; green) shows a cytosolic localisation in the cell after treatment with 10 nM of BFA for three hours, while DAPI stains the *Naegleria* nucleus and mitochondrial DNA. The image is a result of a 3D rendering of 32 individual, 0.29 μm thick sections, overlapping with a final representative thickness of 9.27 μm. Different angles of the same image can been found in **Supplementary Figure 4a**. **e.** Cellular localisation of COPI in two *N. gruberi* cells. Chicken anti-*N.gruberi* COPB antiserum (1:250; green) shows a tubular localisation in the cells after treatment with 1 μM of BFA for three hours, while DAPI stains the *Naegleria* nuclei and mitochondrial DNA. The image is a result of a 3D rendering of 26 individual, 0.29 μm thick sections, overlapping with a final representative thickness of 7.53 μm. Different angles of the same image can been found in **Supplementary Figure 4b**.

Finally, to investigate the subcellular localisation of *Ng*COPB even further, we utilized immune-gold electron microscopy (**Figure 5, Supplementary Figure 6**), which showed with high confidence localisation of this protein in distinct membrane organelles (1 - 4 μm in length) as opposed to the cytosol, nucleus, larger membrane organelles and membrane vesicles.

**Figure 5:**
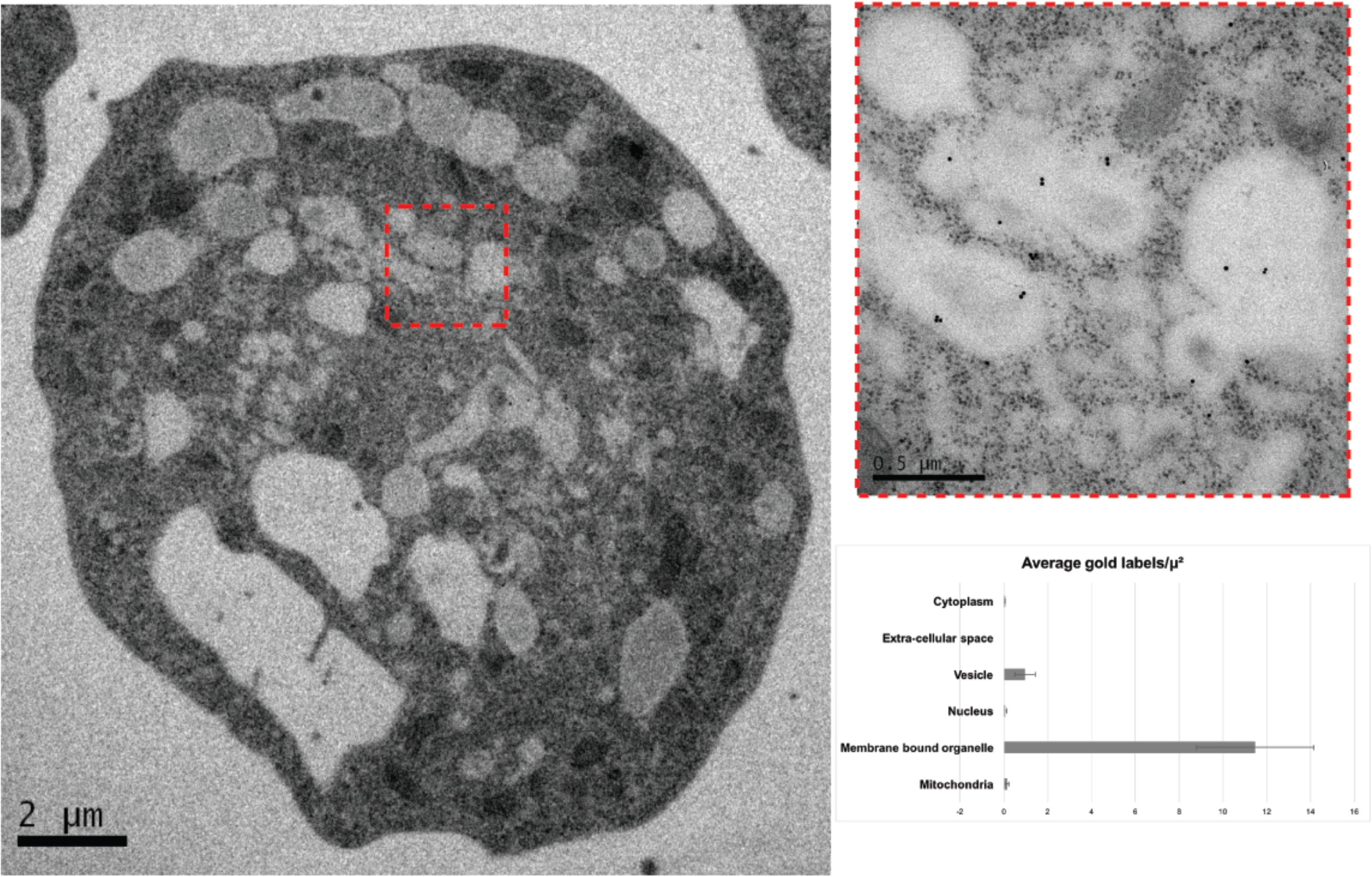
Iiinmino-gold localisation of COPI in *Naegleria gruberi* cells. Immuno-gold localisation of COPI in *Naegleria gruberi* cell by transmission electron microscopy shows localisation associated with membrane bound organelles. The inset shows a higher-magnification image of the organelles. Four additional images can be found in **Supplementary Figure 6**. The graph demonstrates the densities of labelling in the different compartments of *N. gruberi* cells suggesting that COPI is mainly localized in the membrane bound organelles of the cell.

## Discussion

In this study, we have employed a comprehensive set of tools to firstly identify the cryptic Golgi organelle in *Naegleria* and then investigate its ultrastructure. Prior to this study, the existence of a Golgi apparatus in *Naegleria* had only been indirectly inferred through its phylogenetic relationship with stacked Golgi-possessing lineages and, slightly more directly shown, by the presence of a moderate list of genes from a genome of single species, which in canonical systems are localised at the Golgi. We therefore, sought to provide an increasingly direct and convincing array of evidence. Additional homology searching within *N. gruberi* allowed the identification of several previously unreported proteins that are characteristically associated with the Golgi apparatus, while transcriptomic analysis of a second species demonstrated that their presence in another *Naegleria* representative *(N. fowleri)* and more importantly that these genes are indeed expressed. The complement of *Naegleria* Golgi genes suggests both *cis*-Golgi and *trans*-Golgi Network (TGN) functions and retrograde pathways from the *cis*-Golgi to ER, as well as multiple routes from the TGN outward. This is consistent with complex Golgi function in *Naegleria* fits well with other reports of vesicle coat complexity (L. K. L. K. Fritz-Laylin et al. 2010; Hirst et al. 2014).

As COPI is a well-established marker for the Golgi, we generated a homologous polyclonal antibody against the entire *Ng*COPB protein and we used it for subsequent localisation studies. COPB is a peripheral membrane protein that translocates to membranes from the cytosol during vesicle formation for the transport of material from *cis*-Golgi to the ER (Orci et al. 1986; Szul & Sztul 2011; Papanikou et al. 2015). In the case of *N. gruberi*, our immunofluorescence microscopy studies have demonstrated discrete tubular localisation that occupies 17% of the total cellular volume (**Supplementary Figure 7**) and does not show the same localisation patterns as DAPI, Sec31 or ER-tracker. Both confocal and immuno-electron microscopy has provided evidence of the association of this protein with membranous structures. The identity of this membrane structures as Golgi homologues was further confirmed with BFA experiments. Brefeldin A acts on the Arf-GEF, GBF1, blocking COPI vesicle formation (Klausner et al. 1992). Exercising both biochemical and microscopical means, we have demonstrated that BFA observations are consistent with the behaviour of Golgi markers in other organisms with unstacked Golgi (e.g. *Plasmodium, Entamoeba* and *Giardia*) upon BFA treatment (Lujan et al. 1995; Manning-Cela et al. 2003; Struck et al. 2005).

Based upon the sum of our data, we conclude that the Golgi in Naegleria takes the form of discrete tubular compartments, which did not exceed 1 μm in diameter and 4 μm in length. These tubules did not appear concentrated in any specific region of the cell but were dispersed throughout.

There are several intriguing avenues for future investigation of *Naegleria* Golgi. Our data addresses only a *cis*-Golgi marker, leaving open the question of TGN organization. We observed the Golgi in the most common life-stage of *Naegleria*, the trophozoite, but the behaviour of the organelle in the flagellate and cyst forms would be worthy of enquiry. Finally, organellar dynamics would be fruitfully investigated using live-cell imaging and higher-resolution microscopy, such as light-sheet technologies. In *Giardia* and yeast, there is evidence of transient communication intermediates (Stefanic et al. 2009; Szul & Sztul 2011), either tubular or vesicular between the compartments. Understanding the mechanism of material transfer between *Naegleria* Golgi will be an exciting challenge. All of these aspects would be greatly facilitated by additional molecular cell biological tools in *Naegleria* that are currently being developed (protocols.io). Given the complexity of the metabolic and cell biological complement encoded in its genome (L. K. Fritz-Laylin et al. 2010), we suggest that *Naegleria* is a promising model organism for comparative eukaryotic cell biology.

In theory, there is an array of forms that a stacked Golgi could take upon an evolutionary reorganization in a lineage. These include a single large (but unstacked) organelle, a vesicular tubular network, a tubular network, or discrete dispersed smaller compartments. While a single perinuclear organelle is reported in *Plasmodium* (Struck et al. 2005), to our knowledge its ultrastructure has not been examined at the EM level. The organelles of *Giardia, Entamoeba*, and *Saccharomyces* are best described as discrete dispersed small compartments. By contrast, the Golgi in microsporidian lineages is reported as either an array of lamellar membranes in the cysts or as a tubular (but not vesicular) network (Takvorian et al. 2013; Beznoussenko et al. 2007). This latter organization is most similar to that which we observe in *Naegleria*. From this diversity of form, it appears that there are no obvious constraints excluding any of the potential organellar morphologies that Golgi may take upon leaving the otherwise nearly ubiquitous stacked organization. It will be exciting to investigate in additional taxa, and using sophisticated cell biological techniques, to further elucidate the ultrastructure of these compartments. This also raises the fundamental question of why stacking is so pervasive amongst eukaryotes, given the demonstration that other morphologies can readily exist.

## Acknowledgements

We would like to thank P. Melancon, M. Leger and members of the Dacks and Tsaousis labs for helpful discussion. This work was primarily funded by a Royal Society International Exchanges grant (2015/R1-IE150049) jointly to ADT and JBD. Additional funding to ADT was provided by a grant from the Gordon and Betty Moore foundation and the BBSRC (BB/M009971/1), which currently supports LY. Work in the Dacks lab was supported by a Discovery Grant (RES0021028) from the Natural Sciences and Engineering Research Council of Canada. EKH was supported by a Vanier Canada Graduate Scholarship and JBD is the Canada Research Chair (Tier II) in Evolutionary Cell Biology. CNM was funded by an award from the School of Biosciences at the University of Kent.

